# Defining Strawberry Uniformity using 3D Imaging and Genetic Mapping

**DOI:** 10.1101/2020.03.01.972190

**Authors:** Bo Li, Helen M. Cockerton, Abigail W. Johnson, Amanda Karlström, Eleftheria Stavridou, Greg Deakin, Richard J. Harrison

## Abstract

Strawberry uniformity is a complex trait, influenced by multiple genetic and environmental components. To complicate matters further, the phenotypic assessment of strawberry uniformity is confounded by the difficulty of quantifying geometric parameters ‘by eye’ and variation between assessors. An in-depth genetic analysis of strawberry uniformity has not been undertaken to date, due to the lack of accurate and objective data. Nonetheless, uniformity remains one of the most important fruit quality selection criteria for the development of a new variety. In this study, a 3D-imaging approach was developed to characterise berry uniformity. We show that circularity of the maximum circumference had the closest predictive relationship with the manual uniformity score. Combining five or six automated metrics provided the best predictive model, indicating that human assessment of uniformity is highly complex. Furthermore, visual assessment of strawberry fruit quality in a multi-parental QTL mapping population has allowed the identification of genetic components controlling uniformity. A “regular shape” QTL was identified and found to be associated with three uniformity metrics. The QTL was present across a wide array of germplasm, indicating a strong candidate for marker-assisted breeding. A greater understanding of berry uniformity has been achieved through the study of the relative impact of automated metrics on human perceived uniformity. Furthermore, the comprehensive definition of strawberry uniformity using 3D imaging tools has allowed precision phenotyping, which has improved the accuracy of trait quantification. This tool has allowed us to illustrate the use of advanced image analysis towards the breeding of greater uniformity in strawberry.

## Introduction

Strawberries (*Fragaria × ananassa*) are not true fruits. The red fleshy pseudocarp of a strawberry is formed from a swollen flower base or receptacle. The true fruits are, in fact, the achenes which develop from a whorl of carpels and together form an aggregate-accessory fruit. The viability of both carpels and pollen play an important role in the resulting uniformity of berries ^1^. Carpel position, density and viability dictate the shape, size and uniformity of a strawberry. Indeed, strawberry breeders have selected for high carpel densities in order to produce larger fruits ^1^. Simple, classical studies which remove all or part of the achenes from undeveloped pseudocarps has led to a cessation in the auxin “swelling signal” in the area beneath each achene and thus uneven fruit development ^2^. In a similar fashion to achene removal, uneven pollination of the carpels, or absence of achene development, are the main causes of uneven pseudocarps ^3^. Uneven successful pollination can be caused by damage to flowers through high temperature, frost or precipitation ^1^. A late frost in spring could lead to carpel and other damage, resulting not only in malformation but also complete lack of strawberry development ^1^. Strawberry flowers have a variable proportion of viable carpels and anthers between flower orders, both within a plant and also between different cultivars ^4,5^. Indeed, primary fruit are more likely to be malformed due to the relatively lower quantities of viable anthers and pollen ^6,7^.

In spite of the environmental factors known to influence uniformity, literature has shown that strawberry uniformity still has a large genetic component and can be improved through breeding^8,9^. Indeed, where breeders have selected for increased uniformity within and among berries, improvements in uniformity were observed over time ^8^. Cultivars have been shown to differ in their susceptibility to misshapen fruit, indicating a significant genetic component controlling uniformity ^1,8,10^. For example, ‘Florida Elyana’ is susceptible to rain damage, disrupting carpel development and thus misshapen fruit leading to lower market value ^9^, similarly ‘Camerosa’ has been noted as a cultivar which is particularly susceptible to misshapen fruit with ~4% of yields lost as a result of misshapes ^10,11^. By contrast, ‘Florida Radiance’ has high marketable yields and does not exhibit a high proportion of misshapen fruits ^9^. Breeders can influence the proportion of uniform strawberries through selecting- be it directly or indirectly- for 1) even allocation of viable carpels across the receptacle within the flower 2) ready access to pollen within flowers and 3) high fertility of carpels ensuring even successful pollination.

Strawberry is an important fruit crop with a global market revenue of 21,171 million USD in 2015 ^12^. Producing visually appealing strawberry fruit is one of the primary objectives in a strawberry breeding program ^13^. Shape uniformity is an essential trait of strawberry fruits due to the direct association with product quality and value ^14^. Increasing the uniformity of berries can increase the proportion of marketable fruit as berry irregularity is one of the primary imperfections leading to culling and reduced marketable yield ^8^.

As there is no well-defined strawberry phenotyping guidelines for fruit uniformity, the current system at NIAB EMR relies on visual assessments, which are subjective and laborious. Unlike morphological traits such as length, volume and colour, which can be accurately measured manually in a low-throughput manner, uniformity assessment is extremely subjective. As there is no quantitative method of generating phenotypic data for uniformity, the genetic determinants of strawberry uniformity are still unknown.

Computer vision has shown great potential to quantify external fruit quality and 2D imaging has been successfully implemented to measure the shape and size of fruits such as strawberries ^15^, apples ^16^, watermelon ^17^, cherries ^18^ and mangos ^19^. Basic shape traits such as length, width, aspect ratio and volume, and more sophisticated traits such as elliptic Fourier descriptors ^20^ have been quantified and used to describe variation in fruit quality. 3D imaging has been successfully used for phenotyping the crop canopy ^21,22^ and root architecture ^23,24^, and a 3D strawberry phenotyping platform has been explored in our previous study ^25^. With the 3D point cloud reconstructed based on the Structure from Motion (SfM) method ^26^, basic size-related parameters have been measured in three dimensions allowing volume estimation with high accuracy ^27^. Compared with shape and size evaluation, uniformity is a multi-dimensional trait, therefore it is not possible to quantify through 2D image analysis with a single viewing angle. The application of 3D image analysis for phenotyping the external qualities of fruit has not been sufficiently explored, and the basic, previously characterised, shape- and size-related parameters are not adequate for understanding uniformity.

Here the application of a 3D phenotyping platform allows us to investigate the genetic basis of strawberry uniformity. The 3D image analysis software leverages the previously developed platform ^25^ in order to define eight new external variables and investigate their importance on manual uniformity assessment. This method was applied to a multi-parental strawberry mapping population in order to quantify the genetic components underpinning strawberry uniformity.

## Materials and methods

### Plant material and experimental set-up

A multi-parental strawberry population was generated through crossing 26 diverse cultivars and breeding lines to create a population of 416 genotypes made up of 26 families each containing 16 individuals (Suppl. Figure 1). Progenitors were selected to represent diversity across multiple fruit quality traits. Twelve replicate runner plants were pinned down from each genotype into 9 cm square pots containing compost. Clonal plants were separated from parental plants and then placed in cold storage (−2°C) until the start of the experiment. Plants were potted into 2 L pots containing coir and fertigated at 1kg L^−1^ (rate: 10 seconds every 45 min) using Vitex Vitafeed (N:P:K, 176:36:255). Blocks were horizontal intersections across the polytunnels. Due to the large scale of the experiment, replicate blocks were set up at three week intervals. A Natupol Koppert bumble bee hive was added into each polytunnel to assist even pollination. Strawberries were picked when ripe into egg boxes. Boxes were labeled with QR codes to assist tracking of genotypes. Strawberry uniformity was scored on a scale from 1 (irregular) to 9 (uniform) with extensive training provided for all assessors. Strawberry shape was allocated into 9 categories: globose, globose-conic, conic, long-conic, bi-conic, conic-wedge, wedge, square and miscellaneous. Manual uniformity scores were recorded in the field book app ^28^, the QR scanning feature allowed quick access to the correct entry form.

### Genotyping and Linkage map

DNA was extracted for each genotype from unopened leaflets using the Qiagen DNeasy plant mini extraction kit. Genotyping was conducted using the Axiom^®^ IStraw35 384HT array ^29^ (i35k). Crosslink was used to generate linkage maps- a program developed specifically for polyploid plant species ^30^. The map orders from 5 populations were combined to make the consensus map as detailed in the study of Harrison et al ^31^. *Fragaria* × *ananassa* chromosome number is denoted by 1-7 and the sub-genome number is represented by A-D as specified in ^32^.

### 3D reconstruction

The 3D imaging platform was a modified version of that developed by He et al. ^25^. Strawberry fruit were placed in the middle of a turntable, on a dark blue holder made by polymeric foam (38 mm × 19 mm × 19 mm; height, length and width). Unlike the previous study, a webcam (Logitech C920, Newark, CA, USA) was fitted at a height of 30 cm and horizontal distance of 25 cm away from the sample. QR codes on containers were scanned through the webcams allowing tracking of berries and automated labeling of image files. The imaging rig was placed inside a photography studio tent with constant LED illumination. The turntable rotated at a frequency of 50 seconds per full turn, and an image was captured every second. Six imaging platforms allowed concurrent imaging of replicate berries. The 3D reconstruction was implemented with Agisoft Photoscan (Agisoft, LLC, St. Petersburg, Russia), and in order to increase the accuracy and processing speed, all images were pre-processed by cropping to a smaller size (400 × 600 pixels). Background subtraction was achieved through arbitrary colour thresholding. The image processing software for webcam control and automated image pre-processing were written in C++ utilising the OpenCV Library ^25,33^.

### Data processing pipeline of phenotypic traits extraction

#### Point cloud preprocessing

In the preprocessing stage (Fig. 1), each point cloud model was converted from the colour space of RGB (Red, Green and Blue) to HSV (Hue, Saturation and Value). Arbitrary thresholding on the hue channel was used to remove the noise introduced in the reconstruction stage. The clean point cloud was translated to the origin of the 3D coordinate system based on the distance between the moment of the point cloud and the origin. By calculating the eigenvector associated with the largest eigenvalue of the coordinates of points, a rotation matrix could be derived to represent the main orientation of the point cloud, which can be used to rotate the point cloud with the main orientation aligned with the z-axis. After rotation, the arbitrary threshold was applied again on the hue channel in order to segment the strawberry body and blue holder from the whole point cloud. The height of the holder was obtained by calculating the difference between the maximum and minimum values of the holder point cloud on the z-axis. As the original coordinate system generated by Structure from Motion (SfM) method has an arbitrary scale, each point cloud model needed to be standardised by the height of the holder, so that the sizes of all point clouds are comparable.

**Figure 1.**
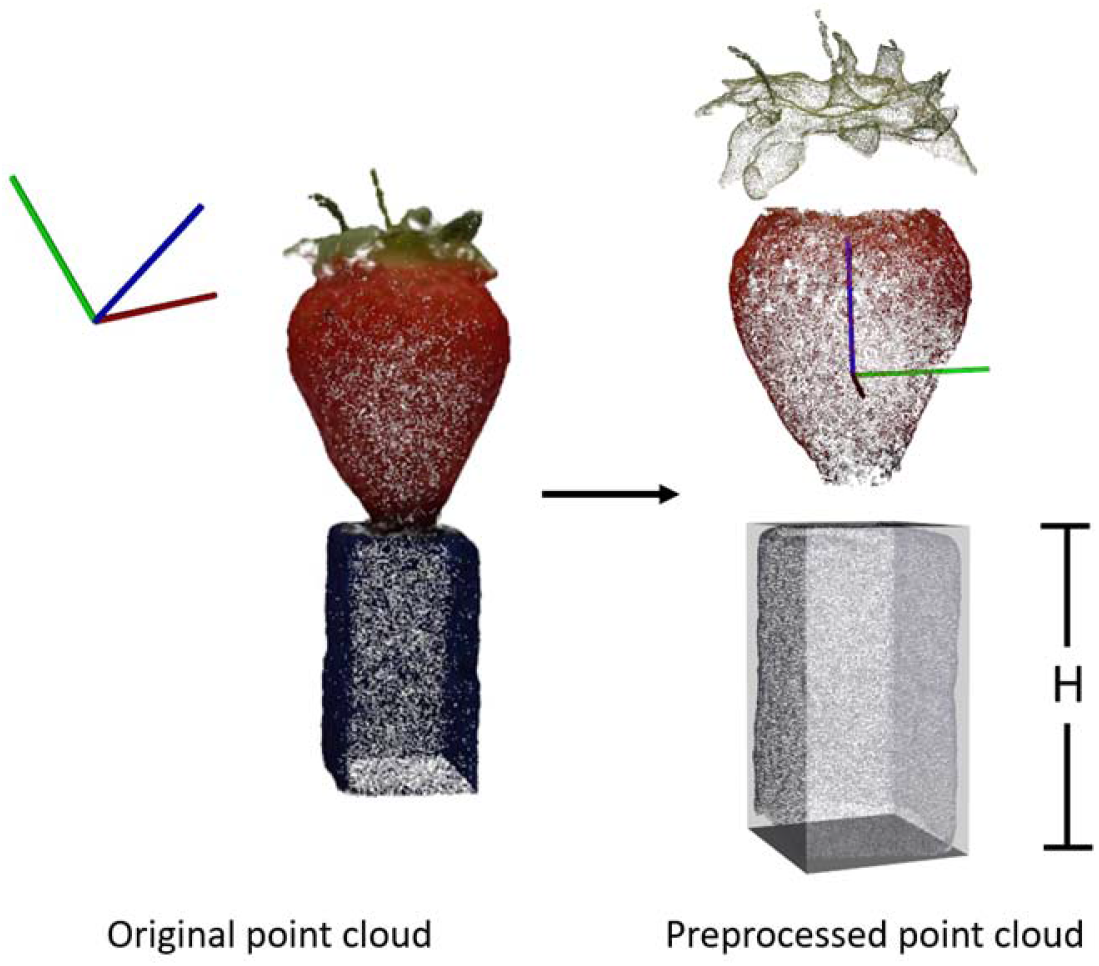
Point cloud pre-processing for strawberry body extraction, translation to origin of xyz coordinate system and size standardisation.

#### Uniformity-related traits measurements

Eight uniformity-related traits were calculated from the point cloud data of strawberry body after preprocessing. These are:

### Coefficient of variation (CV) of side view areas (CV_A) and the ratio between maximum and minimum side view area (Max_A/Min_A)

All side views should be identical in a perfectly uniform strawberry. In order to eliminate the heterogeneity introduced from the calyx and the holder, only the points within the middle 50% of the body height of each point cloud were retained for analysis (Fig. 2). In order to understand the heterogeneity of different side views of a point cloud, each point cloud was rotated along the z-axis by 3.6° for 99 rotations, and after each rotation, the side view of the point cloud was projected onto the x-z plane in 2D (labelled in white). A convex hull was fitted to each projected image and the contour area was calculated. For area metrics, two traits were obtained; the CV of side view areas (CV_A) and the ratio between the maximum and minimum area (Max_A/Min_A). An ideal uniform strawberry will have a value of zero for CV_A and one for Max_A/Min_A.

**Figure 2.**
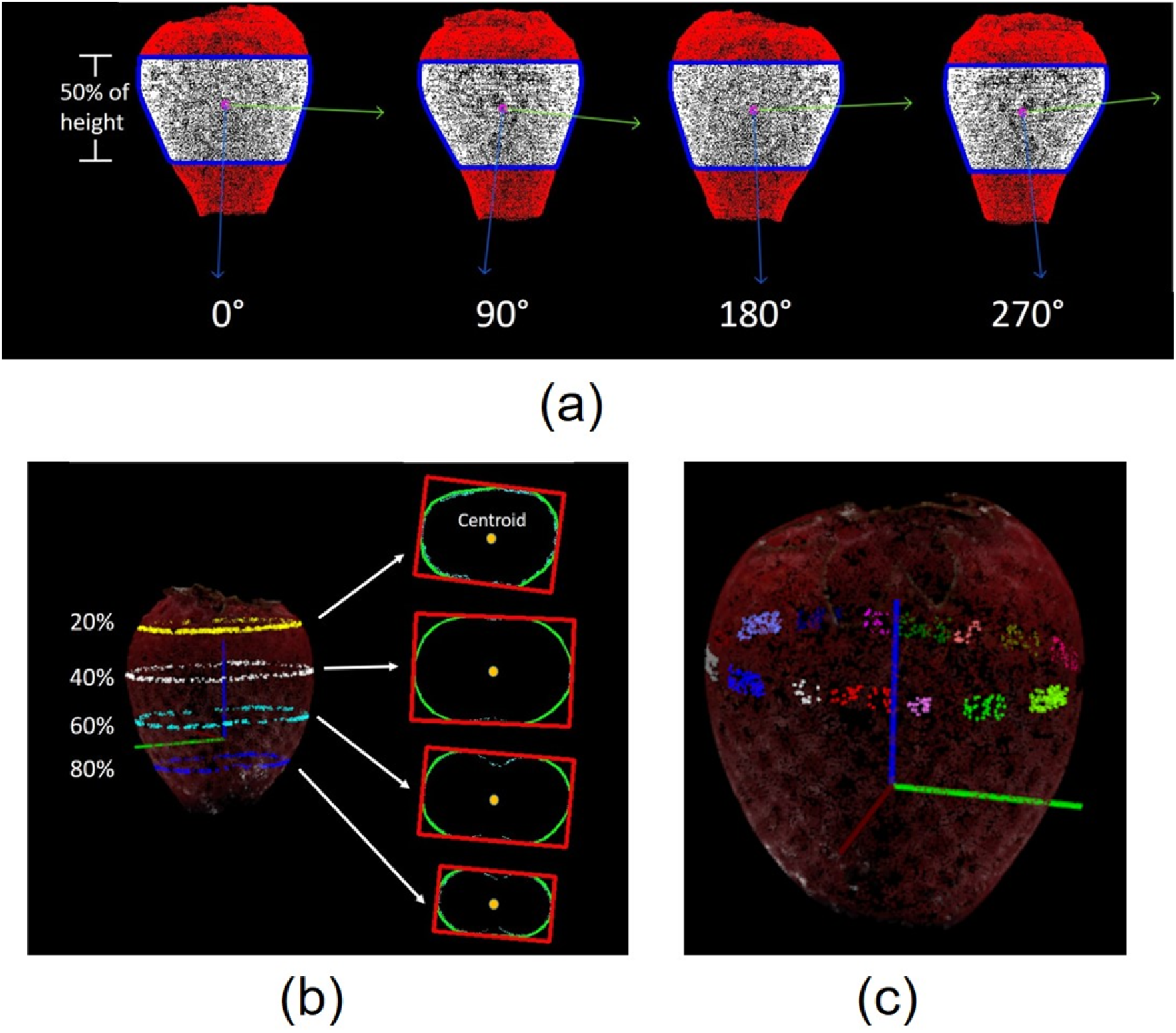
Side view of strawberry body for the CV measurement of the area and principal orientations. Convex hulls are outlined in blue, and blue and green arrows indicate the principal orientations (a). Extraction of example slice images horizontal to x-y plane at the height of 20%, 40%, 60% and 80% of the total height. A minimum bounding box is fitted to each slice image (b). Sixteen patches of points labelled in different colours for curvature estimation (c).

### CV of principal orientations (CV_D)

The major eigenvector indicating the main orientation was calculated by principal component analysis (PCA) for all 100 side view projected images, and the heterogeneity of the orientations of the projected images was quantified by calculating the CV of angles of the main orientation. Like CV_A, a perfectly uniform strawberry will have a value of zero for CV_D.

### Aspect ratio of the minimum bounding box (L/W)

A lateral slice image was obtained by identifying the intersection between the plane in parallel with the x-y plane and point cloud (Fig. 2b). Based on the values on the z-axis, 100 evenly spaced slice images were obtained. The slice image with the largest contour was obtained by calculating the contour area of the convex hulls for all slice images. The main orientation of the contour was indicated by the major eigenvector of the PCA and the minimum bounding box was fitted to the slice images. The ratio between the length and width of the largest bounding box was derived and the ratio should be one for a perfectly uniform fruit.

### Circularity of the maximum circumference (CIR)

Visually, the circularities of the contours in horizontal slice images are high if the strawberry is uniform. Circularity (CIR) was calculated as previously described ^34^:

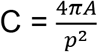

Where *A* and *p* are the area and perimeter of the convex hull respectively. For each point cloud, the circularity was calculated for the slice image with largest contour area.

### Straightness of centre axis (STR)

The coordinates of the centroids for each horizontal slice image can be located by calculating the moment of the contour. The centroids can be connected as a straight line for a uniform strawberry. The centroids were calculated for all the slice images within the middle 80% of the body height, and the straightness of the central axis was characterised by:

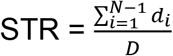

Where N (N = 80) is the number of slice images used for the analysis, d_i_ is the Euclidean distance between neighbouring slice images, and D is the Euclidean distance between the centroids of the top and bottom slice images.

### CV of curvature and the ratio between maximum and minimum curvatures (Max_C/Min_C)

The principal curvature can be calculated for each point in the point cloud, which describes how much the curve deviates from a straight line at this point. It can be imagined that the 3D curve can be sliced orthogonally around the direction of normal in to 2D curve, and the maximum curvature k_1_ and minimum curvatures k_2_ are the two principal curvatures for the 3D curve ^35^. The average curvature k, which is defined as the mean value of the magnitudes of principal curvatures in the two main directions was applied to quantify the curvature for a given point.

As the curvature measurement is sensitive to noise, the point cloud surface of strawberry body was first smoothed by using Moving Least Squares (MLS) method ^36^, which could reconstruct a smooth surface from the noisy point cloud. Sixteen patches of the points were selected evenly from the points forming the largest slice in parallel with x-y plane (Fig. 2c). For each patch, the first half largest curvatures were averaged and used to represent the curvature of the patch. With the curvatures of all 16 patches, the CV of curvature (CV_C) and the ratio between maximum and minimum curvatures (Max_C/Min_C) were calculated.

#### Statistical analysis

##### Ordinal regression

Statistical analysis was performed using R (version 3.5.1) and the Genstat statistical package (Version 13.0, VSN International Ltd. England). Differences in uniformity traits within each shape groups were distinguished using ANOVA and Tukey post-hoc test. Pearson coefficients of correlation were calculated between all proposed uniformity-related traits. As the group labels are ordinal dependent variables, ordinal regression was used to evaluate the performances of all traits ^37^. Model fit was ascertained by using selection criterion values based on the Akaike Information Criterion (AIC) and the Bayesian Information Criterion (BIC). In general, a better model fit generates lower values for both AIC and BIC ^38^. In order to identify the optimal variable combination related to manual assessment, stepwise AIC and BIC methods were applied ^39^. The most significant variable was identified by comparing the criterion values of all models. Other variables were added successively and retained if the model fit was improved.

##### Genetic Analysis

The best linear unbiased estimate (BLUE) was calculated for all genotypes in order to correct for the influence of assessor, data and block. Linear mixed-effects models were generated for each phenotypic trait with and without covariates. Grand scores for each genotype were calculated using mixed models to account for significant covariates.

##### Composite interval mapping

Multiparental QTL mapping was conducted in R using package “mppR” ^40^. A permutation test determined the significance threshold ^41^. A two-step QTL analysis was implemented: the selection of cofactors was achieved through Simple Interval Mapping (SIM) proceeded by a multi-QTL model search using composite interval mapping (CIM) ^42,43^. As a multi-parent population CIM works on parent relationships. Therefore the ‘CPEM0162’ × ‘Rumba’ cross was removed as it is not directly related through the parental cultivar network. All other crosses were interrelated and formed a single network (Sup. Figure 1).

## Results

### Characterisation of uniformity-related traits

All the uniformity-related traits were calculated based on the point cloud, the mean values and the standard errors for each visual uniformity class are presented in Figure 3. ANOVA results showed that significant differences were observed between uniformity classes for all traits (*p* < 0.001). The Pearson’s linear correlation coefficients were calculated between all traits, and strong correlations were found amongst Max_A/Min_A, L/W and CIR (Table 1).

**Table 1.**
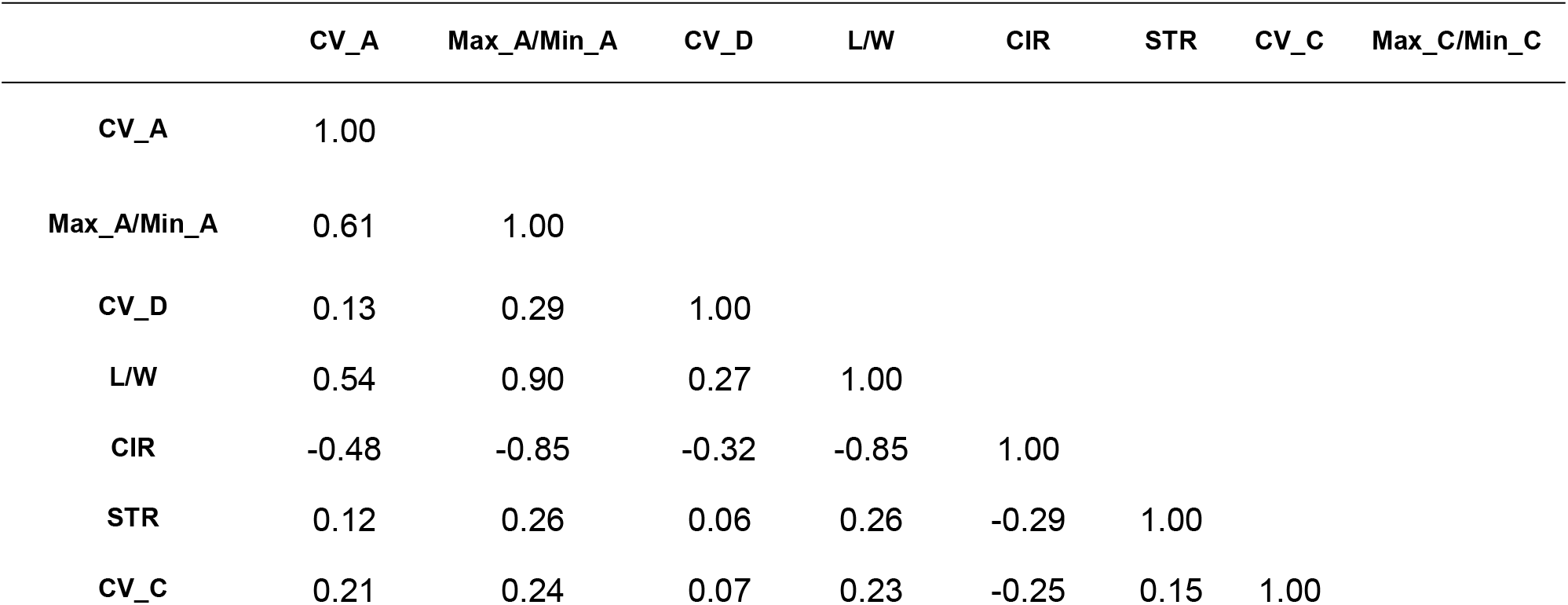
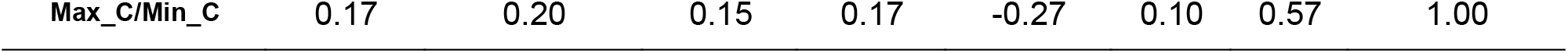
Pearson’s linear correlation coefficients among all uniformity-related traits. All values are significant at *p* < 0.05 level.

|  | CV_A | Max_A/Min_A | CV_D | L/W | CIR | STR | CV_C | Max_C/Min_C |
| --- | --- | --- | --- | --- | --- | --- | --- | --- |
| CV_A | 1.00 |  |  |  |  |  |  |  |
| Max_A/Min_A | 0.61 | 1.00 |  |  |  |  |  |  |
| CV_D | 0.13 | 0.29 | 1.00 |  |  |  |  |  |
| L/W | 0.54 | 0.90 | 0.27 | 1.00 |  |  |  |  |
| CIR | -0.48 | -0.85 | -0.32 | -0.85 | 1.00 |  |  |  |
| STR | 0.12 | 0.26 | 0.06 | 0.26 | -0.29 | 1.00 |  |  |
| CV_C | 0.21 | 0.24 | 0.07 | 0.23 | -0.25 | 0.15 | 1.00 |  |

**Figure 3.**
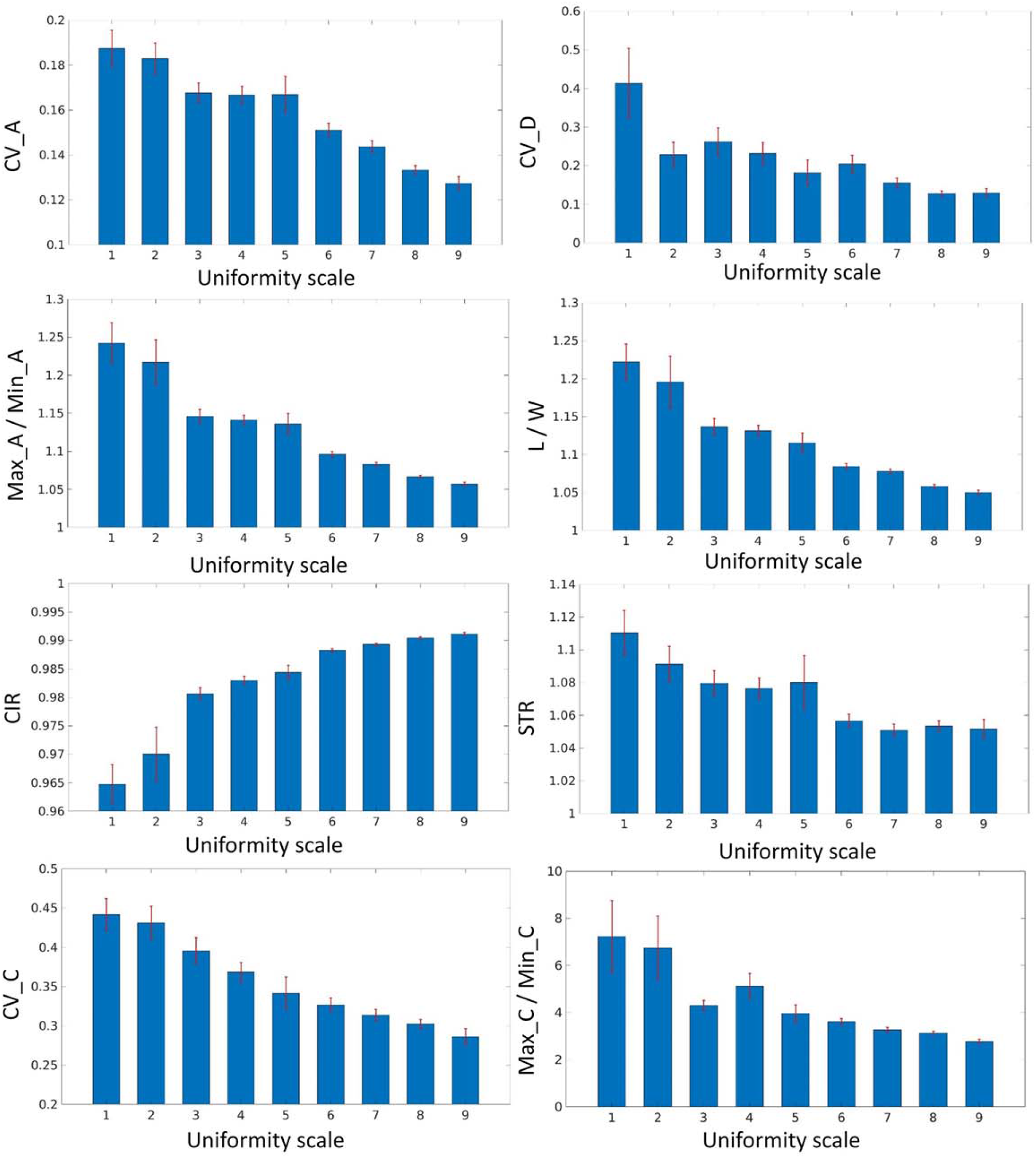
Mean value and standard error of calculated uniformity-related traits by the newly developed 3D image analysis software against defined uniformity scale based on manual assessment.

Ordinal regression models were constructed for all variables and each variable independently. L/W was not significant due to the high correlation with other variables and CIR showed the best model fit with the lowest AIC and BIC values (Table 2). New variables were added sequentially to the model until no further improvement of the criterion value was observed. The AIC and BIC based stepwise selection methods showed inconsistent results (Table 3). The AIC based method showed the optimal criterion value with all variables except L/W and Max_C/Min_C, but BIC based method showed that STR could not improve the model fit.

**Table 2.**
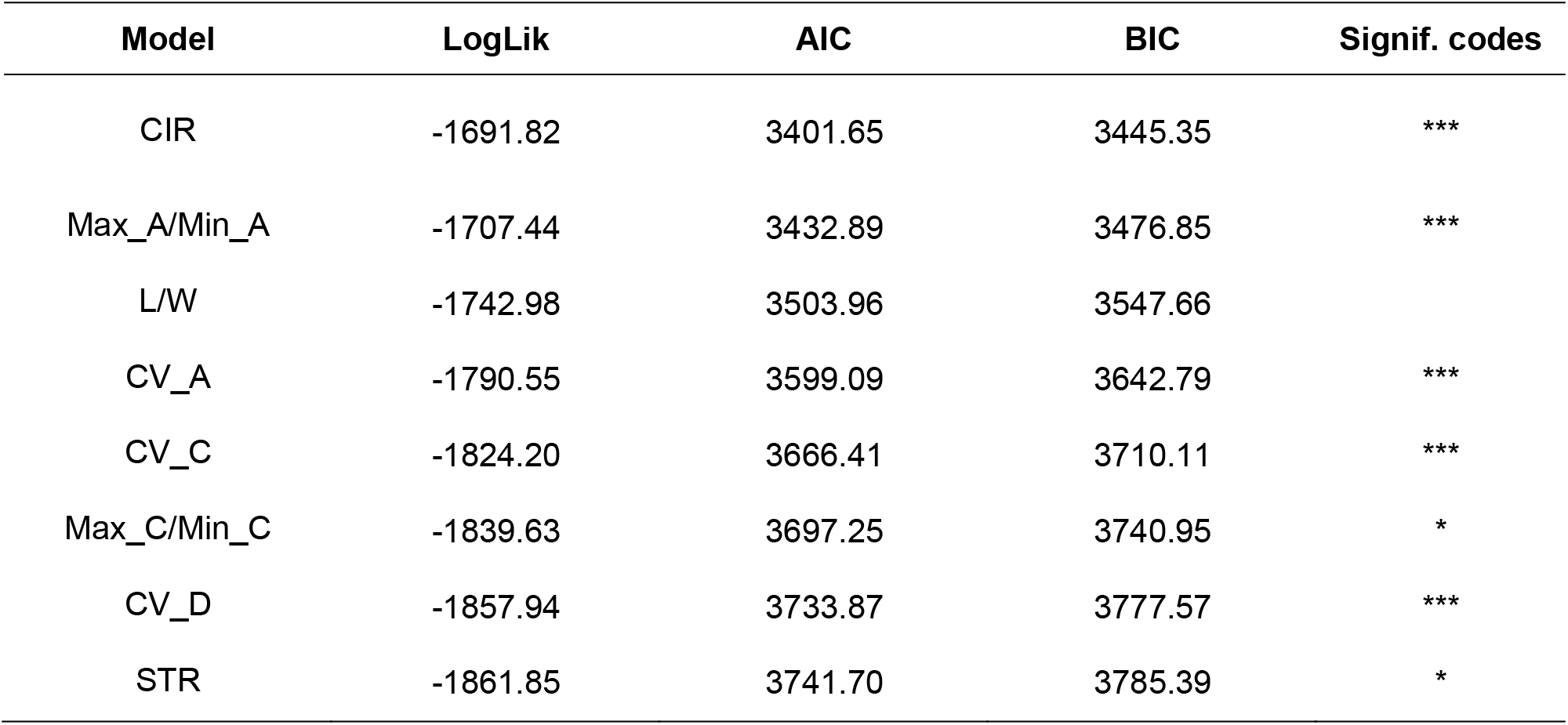
Summary of individual ordinal models and variable significance of ordinal model with all variables, towards prediction of manual assessment uniformity scores. * p < 0.05, ** p < 0.01, *** p < 0.001.

| Model | LogLik | AIC | BIC | Signif. codes |
| --- | --- | --- | --- | --- |
| CIR | -1691.82 | 3401.65 | 3445.35 | *** |
| Max_A/Min_A | -1707.44 | 3432.89 | 3476.85 | *** |
| L/W | -1742.98 | 3503.96 | 3547.66 |  |
| CV_A | -1790.55 | 3599.09 | 3642.79 | *** |
| CV_C | -1824.20 | 3666.41 | 3710.11 | *** |
| Max_C/Min_C | -1839.63 | 3697.25 | 3740.95 | * |
| CV_D | -1857.94 | 3733.87 | 3777.57 | *** |
| STR | -1861.85 | 3741.70 | 3785.39 | * |

**Table 3.** Model comparison values for uniformity metrics, towards prediction of manual assessment uniformity scores based on AIC and BIC.

| Model | 1 | 2 | 3 | 4 | 5 | 6 | 7 | 8 |
| --- | --- | --- | --- | --- | --- | --- | --- | --- |
| CIR | x | x | x | x | x | x | x | x |
| Max_A/Min_A |  | x | x | x | x | x | x | x |
| L/W |  |  | x |  |  |  |  |  |
| CV_A |  |  |  | x | x | x | x | x |
| CV_C |  |  |  |  | x | x | x | x |
| Max_C/Min_C |  |  |  |  |  | x |  |  |
| CV_D |  |  |  |  |  |  | x | x |
| STR |  |  |  |  |  |  |  | x |
| **AIC** | 3401.65 | 3374.40 | 3374.29 | 3362.57 | 3302.01 | 3299.53 | 3291.87 | 3288.40 |
| **BIC** | 3445.35 | 3422.95 | 3427.70 | 3415.97 | 3360.28 | 3362.65 | 3354.99 | 3356.38 |

### The influence of Shape on uniformity

The shape of a strawberry influences the uniformity trait score. Bi-conic strawberries were seen to have high uniformity based on the area overlap measures (CV_A & Max_A/Min_A), L/W and CIR scores indicating bi-conic strawberries have consistently circular horizontal transects at the mid point. Whereas for curvature uniformity measures (CV_C & Max_C/Min_C) globose fruit are the most uniform and miscellaneous fruit the least (Data not shown). Both the manual uniformity score and CIR could discriminate miscellaneous shapes from the other shape categories (Figure 4).

**Figure 4.**
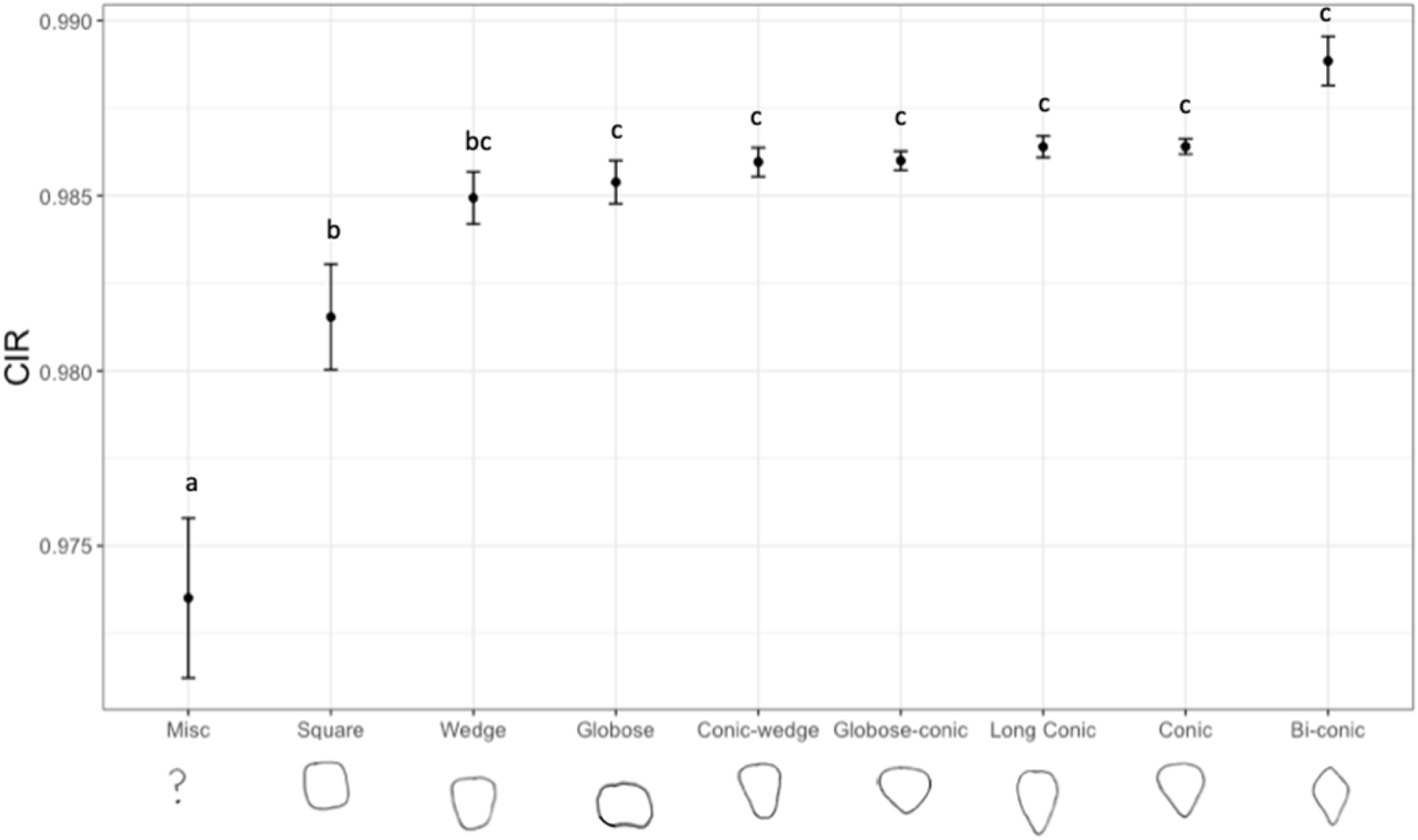
CIR scores for each manually classified strawberry shape category. Letters denote significant differences between categories. Error bars are standard errors of the mean.

### QTL identification

A total of 28 QTL were found to be associated with uniformity traits (Table 4). Of which 25 were detected in more than one progenitor (Figure 5). Five focal SNP’s, on chromosome 2B and 5D were found to represent more than one trait (Table 4, Figure 6). The same focal SNP AX.166521303 was identified as important region in Max_A/Min_A, CV_A and CIR uniformity traits. Global adjusted R^2^ values for linear models were between 5.07 and 32.15 indicating the proportion of variation explained by identified QTLs (Table 5). All uniformity traits apart from CV_A were significantly affected by date of picking. CV_D had the largest broad sense heritability score of 38.4 (Table 5).

**Table 5.** Broad sense heritability scores for each automated uniformity trait, the influence of block and date of assessment on the trait measured. The number of QTL and the coefficient of determination associated with combined QTL.

| Trait | Heritability | Significance of Block | Significance of Date | GxE | Number of QTL | R2 | R2 adj |
| --- | --- | --- | --- | --- | --- | --- | --- |
| STR | 21.70 | *** | *** | *** | 1 | 5.07 | 4.63 |
| CIR | 22.00 | *** | *** | *** | 5 | 21.2 | 19.3 |
| CV_D | 38.40 | *** | *** | *** | 3 | 9.35 | 8.05 |
| CV_A | 19.95 | NS | NS | *** | 2 | 12.13 | 11.29 |
| CV_C | 24.63 | *** | *** | *** | 1 | 6.41 | 5.97 |
| L/W | 19.20 | *** | ** | *** | 6 | 24.77 | 22.59 |
| Max_C/Min_C | 16.70 | NS | * | NS | 7 | 32.15 | 29.85 |
| Max_A/Min_A | 22.70 | ** | *** | *** | 4 | 23.1 | 21.62 |
| Uniformity | 8.00 | NS | *** | *** | 1 | 6.9 | 6.46 |

**Figure 5.**
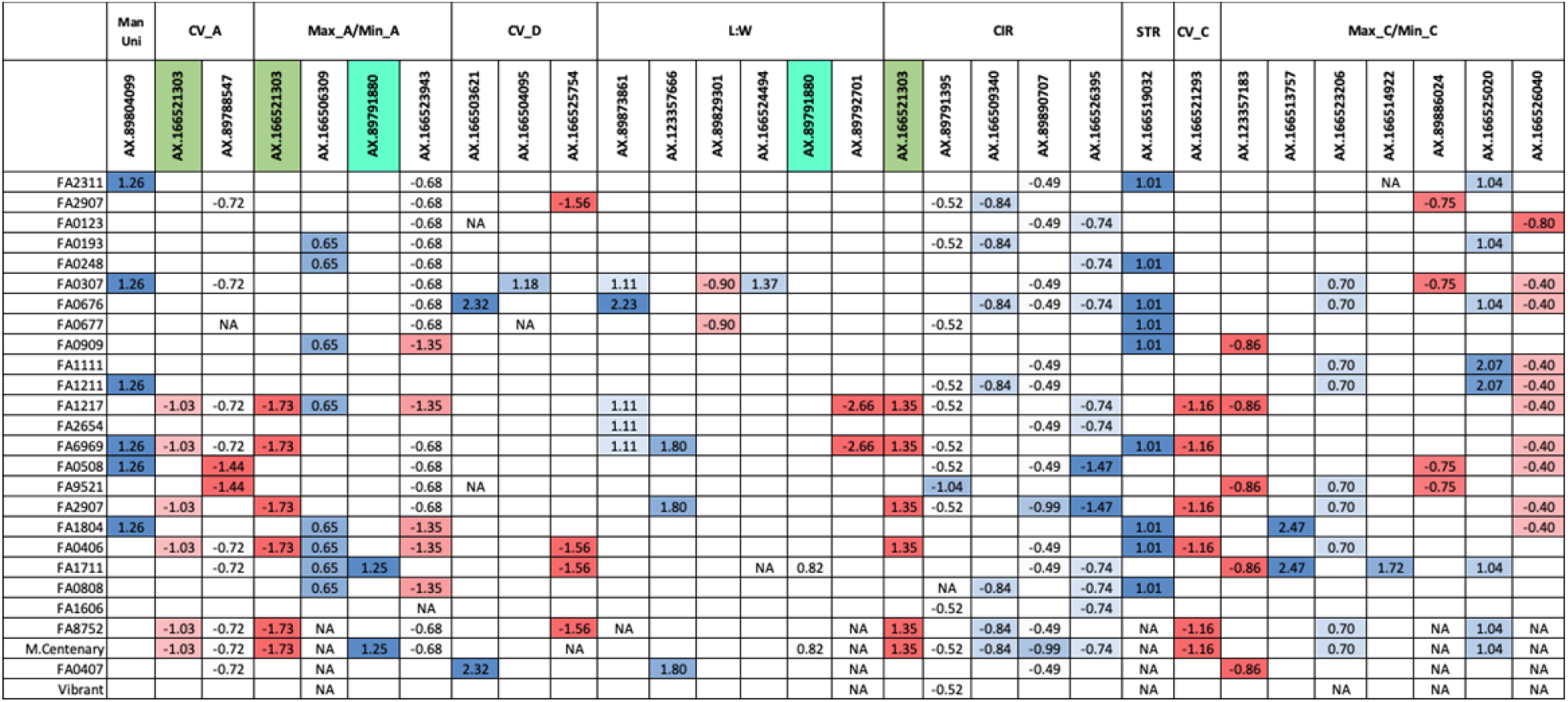
Effect sizes associated with each QTL in each of the 26 progenitors; blue colour is associated with lower uniformity and red colour is associated with higher uniformity.

**Figure 6.**
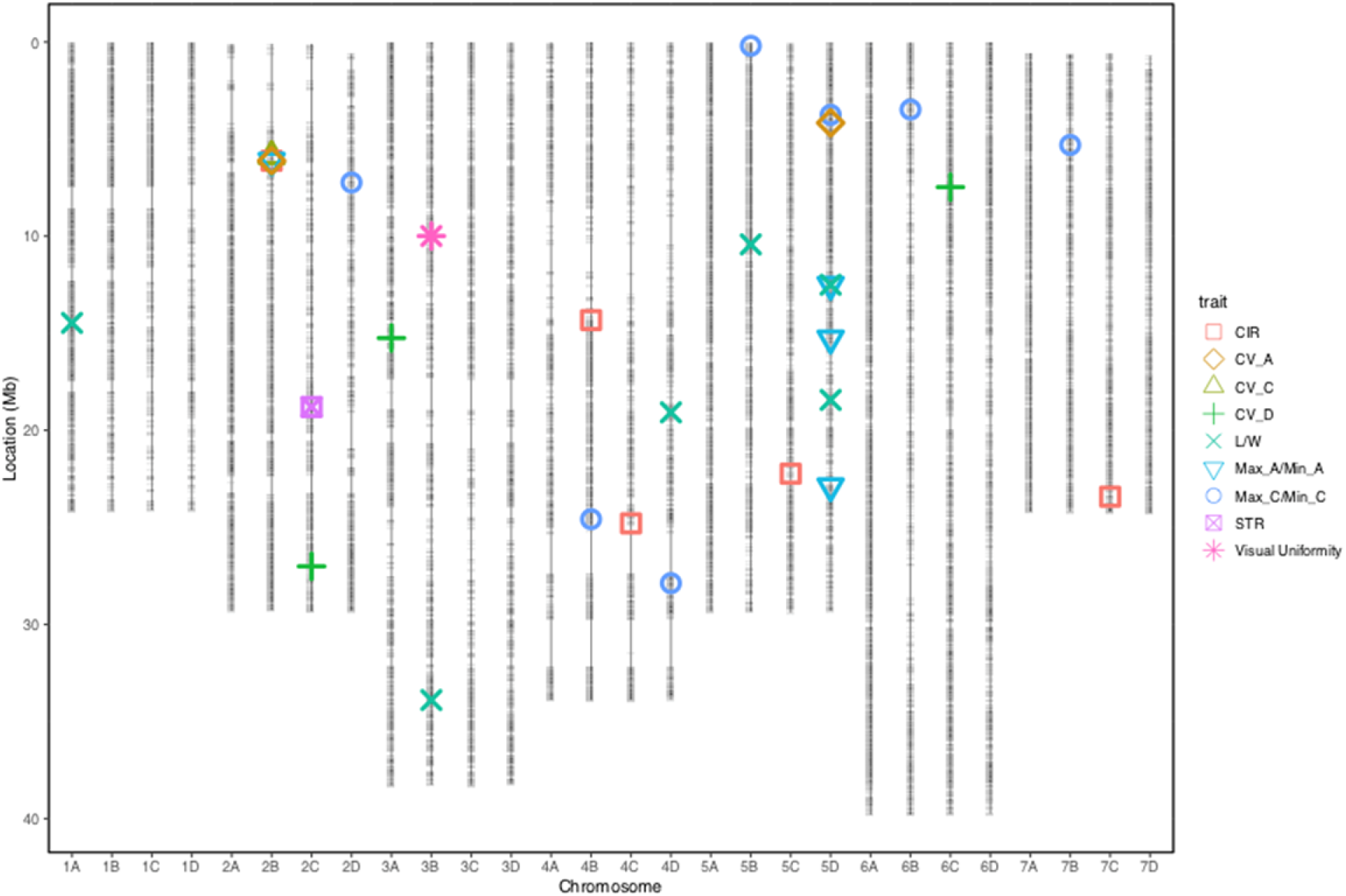
Location of QTL on the octoploid consensus map scaled to the *Fragaria vesca* ‘version four’ genome. Horizontal grey lines represent istraw 35k axiom array markers

## Discussion

We report for the first time a robust method to measure strawberry uniformity and apply this technique to generate genetic markers for uniformity traits. Several studies have attempted to quantify strawberry fruit shape using 2D images with neural networks ^44^, 3D imaging ^45^ and by machine learning^15^. However, none of these studies investigated berry uniformity. Unlike the aforementioned studies, who measure a relatively small number of genotypes intensively, we have implemented a high throughput imaging platform across a large population to facilitate genetic analysis of the trait. Although strawberry shape has received greater attention in the literature, berry uniformity is a more important trait for a breeder to improve (Personal communication, Abigail Johnson).

In current strawberry breeding practice, there is no widely accepted criteria for quantifying uniformity due to the difficulty of defining a multidimensional trait. Here, the manual strawberry uniformity scale has been designed by NIAB EMR breeders. As such, the absence of a straightforward definition, has meant that it has not been possible to study the genetic components controlling strawberry uniformity in the past. To overcome this, we have used 3D image analysis to define the parameters underlying a breeder’s perception of strawberry uniformity. The original 3D strawberry phenotyping system ^25^ could accurately measure basic size-related traits. In this study, the point cloud analysis software was further developed to quantify strawberry uniformity through eight proposed metrics. By comparing with the manual scale, the image processing pipeline has demonstrated an objective method of characterising strawberry uniformity components.

### Quantifying berry uniformity

Circularity of the maximum circumference (CIR) of strawberries showed the best predictive ability for manual uniformity scores based on the ordinal regression model fit, when studying individual variables alone. A completely misshapen fruit with a severely undulating fruit surface will score a value of 1 for manual assessments, and these completely misshapen fruits were the easiest category to identify by eye, as they were clearly distinct from regular shapes. A low CIR value appears to represent the undulating misshapen and “miscellaneous” fruit (Figure 4 & 6). Miscellaneous berries are the most undesirable fruit shape category therefore it is highly beneficial to select against them. When multiple traits are combined to describe uniformity, the best fitting model required the combination of CIR, CV_A and Max_A/Min_A, CV_D and CV_C. The five factors required for optimal model construction indicate that there are multiple uniformity components influencing the manual uniformity score.

### Misshapen fruit QTL

One of the QTL represented by the focal marker AX.166521303 on chromosome 2B was found to be associated with CIR, this QTL was also associated with CV_A and Max_A/Min_A, each of which were found in the best fitting model used to describe the manual uniformity score. The focal SNP AX.166521303 was found to be present and significant in six progenitors and had an effect size of 9.14% on CIR. Therefore, this marker is a good candidate for marker assisted breeding in selection against completely mis-shapen and irregular strawberries. Furthermore, this work has highlighted a region of interest for further study to pinpoint the causative allele associated with reduced uniformity. Dissecting the contribution of genetic and environmental components believed to underpin strawberry uniformity; susceptibility to heat stress, carpel and pollen viability, achene position, size and distribution^1^ may help to further elucidate the mechanism of uniformity segregating in the multiparental population.

### Uniformity trait selection

The trait L/W shows little improvement on the overall combined trait model fit due to the high correlation with other traits including Max_A/Min_A and CIR, but it was still a good predictor of uniformity based on the model fit when studying individual variables alone. AIC and BIC based stepwise feature selection showed disagreement on the selection of the STR parameter. The difference between calculating AIC and BIC is that AIC does not account for the sample size, so when sample number is large, BIC applies larger penalty for complex models and leads to a simpler model ^46^. However, this study does not aim to identify the optimal feature combination to develop prediction model related to manual uniformity evaluation, but develop a new image based quantification to replace the manual scale, because the ground-truth data are subjective and as such any large bias can reduce the robustness of model development. Moreover, the manual scale cannot be considered a comprehensive assessment as the parameter STR cannot be visually evaluated by eye. However, it must be said that if a trait cannot be detected by the human eye, then it is not a valuable trait for a strawberry breeder to select upon.

### Limitations of the system

The 3D point cloud analysis software is independent of the imaging acquisition system, and the uniformity-related traits can be extracted automatically in a high-throughput manner. However, the imaging collection throughput was 50 seconds per fruit and the 3D reconstruction has to be performed separately, which limits use to pre breeding experiments as opposed to use as a breeders tool. Due to the occlusion from the viewing angles, the strawberry nose cannot be fully reconstructed especially for globose shaped fruit, which decreases the accuracy of STR measurements and also limits the study on automated shape classification. To increase the throughput and accuracy of 3D phenotyping, it is necessary to further develop the hardware with multiple cameras to allow more viewing angles or a structured light based imaging system with a robotic arm, and also integrate the hardware driver with 3D reconstruction software. The current point cloud image analysis software was able to characterise many key external traits which are important for strawberry breeding, however, the measurement of other parameters such as achene density must be investigated through an improved phenotyping platform in future studies.

### Genetic Control of Fruit Quality

Papers detailing strawberry fruit quality QTL report genetic alleles associated with multiple fruit quality traits including fresh weight, metabolites, external colour and firmness ^29,47,48^, however, there are currently no papers which report QTL associated with strawberry uniformity. Here, we provide a phenotyping platform which has facilitated the assessment of the genetic components underlying strawberry uniformity for the first time. The use of a multi-parental population has allowed the study of a diverse set of germplasm and has ensured that resulting QTL to have a greater relevance for breeders when compared to alleles identified in bi-parental studies. Overall, 25 of the QTL were found to have an effect on uniformity in more than one of the 26 progenitors indicating that there has been limited linkage decay between the causal allele and marker, and that the relationships have been maintained across generations. Furthermore, the QTL on chromosome 2B was observed three times across different uniformity traits, such traits are only partially correlated and thus describe discrete components, as such this allele can be seen to play a major role in uniformity.

### Genetic control of strawberry fruit shape

Unlike uniformity the mechanism controlling fruit shape has been studied extensively in the wild strawberry; *Fragaria vesca* and may act as a surrogate model for the cultivated octoploid strawberry *Fragaria* × *ananassa*. In *F. vesca*, fruit shape is primarily controlled by phytohormones ^49,50^. Auxin increases the width of fruit and by contrast gibberellic acid (GA) increases the length of a strawberry whereas Abscisic acid (ABA) down regulates both Auxin and GA and thus reduces fruit expansion ^49,50^. GA deficient Vesca mutants were found to have a “short” or globose berry shape, which, through the application of GA to the berry, could be restored to result in a “long” or long conic fruit shape ^50^.

It is clear that breeders wish to select for greater berry uniformity however the confounding relationship between shape and uniformity must also be considered. For example, square, wedge and wedge-conic strawberries may have high 2D symmetry but not 3D symmetry. UK breeders primarily aim to select for conic or long conic fruit whereas globose, square, wedge and miscellaneous berries are classified as undesirable and biconic, globose-conic and conic-wedge fruit are seen as acceptable shapes (Personal Communication, Abigail Johnson). Here we provide an objective measure (CIR) that can be used to discriminate the least desirable berries - miscellaneous or misshapen berries and select for regular fruit shapes.

### Heritability of Uniformity

Broadsense heritability scores were between 16.70 and 38.40 for automated uniformity metrics indicating a greater genetic component than that associated with manual uniformity (8.00). These values indicate the proportion of variation segregating in the study population, however improvement in the heritability may also be caused, in part, by more accurate phenotypic measurements. In particular, high heritability was observed for CV_D which indicates the angle of a strawberry related to whorl of carpels (Figure 4) is under strong genetic control. Date of picking was seen to have a significant impact on all uniformity metrics apart from CV_D which had a large genetic component. The high significance of date indicates the developmental environmental conditions has a significant impact on strawberry uniformity. Extreme temperatures were observed during the experiment which may have caused the significance of date. All traits apart from CV_C showed a significant genotype by environment interaction indicating that genotypes were responding differently to heat stress. Misshapen fruit have been found to have a greater proportion of small underdeveloped achenes following exposure extreme temperatures during embryo development ^51,52^.

Here we provide a comprehensive dissection of the traits underlying strawberry uniformity and show that the visual perception of a strawberry can be represented by 5 metrics. The generation of an objective measure of uniformity has allowed the assessments of genetic components in a multi-parental breeding population. We show uniformity has a strong genetic component that can be improved by breeding and identify genetic components controlling uniformity that are present across a wide array of germplasm.

## Availability of data, materials and methods

The software developed and datasets generated and analysed during the current study are available from the corresponding author on reasonable request.

## Acknowledgements

The authors acknowledge project partners Soloberry, Sainsburys, Botanicoir and Agrovista for their involvement and support of the project. The authors acknowledge Robert Vickerstaff for generating the octoploid consensus map as part of other projects and Dr Beatrice Denoyes, INRA and Dr Amparo Monfort, CRAG for granting the use of their informative markers in the production of the strawberry consensus linkage map. This work was supported by grants from the Biotechnology and Biological Sciences Research Council (BBSRC) BB/M01200X/2, BB/P005039/1 and Innovate UK project 101914.

## Authors’ contributions

BL – Analysed imaging data

AJ, HMC, ES, BL, RJH - Conceived and designed experiments

HMC – Conducted quantitative genetics analysis

AJ, HMC, AK - Performed experiments

AK - Performed genotyping

GD - Ordinal regression analysis

BL, HMC & RJH wrote the manuscript with contributions from all authors.

BL and HMC equally contributed to the manuscript.

## List of Abbreviations

CIM: composite interval mapping
CIR: Circularity
CV_A: Coefficient of Variation of side view areas
CV_C: Coefficient of Variation of curvatures
CV_D: Coefficient of Variation of principal orientations
i35k: Istraw35 Affymetrix chip
L/W: Aspect ratio of the minimum bounding box
Max_A/Min_A: ratio between maximum and minimum side view areas
Max_C/Min_C: ratio between maximum and minimum curvatures
MedR: Median number of roots
QTL: Quantitative Trait Loci
QR: Quick Response
RGB: Red Green Blue
SNP: Single Nucleotide Polymorphism
STR: Straightness of centre axis

---

**Supplementary Table 4.** Focal SNPs representing strawberry uniformity QTL. The position of QTL is reported in Mb as scaled to the vesca version 4 genome

| Marker Names | Chromosome | Pos Mb | $\log^{10} p$<br>value | R<sup>2</sup> | Trait |
| --- | --- | --- | --- | --- | --- |
| AX.89804099 | 3B | 96.3 | 3.29 | 6.9 | Visual Uniformity |
| AX.123357183 | 2D | 63.1 | 4.07 | 3.81 | Max_C/Min_C |
| AX.166513757 | 4B | 239.6 | 3.03 | 5.05 | Max_C/Min_C |
| AX.166523206 | 4D | 272.0 | 3.89 | 4.03 | Max_C/Min_C |
| AX.166514922 | 5D | 1.8 | 3.02 | 9.9 | Max_C/Min_C |
| AX.89886024 | 5D | 36.8 | 3.32 | 2.87 | Max_C/Min_C |
| AX.166525020 | 6B | 34.4 | 3.49 | 7.59 | Max_C/Min_C |
| AX.166526040 | 7B | 47.5 | 3.07 | 2.71 | Max_C/Min_C |
| AX.166521293 | 2B | 60.7 | 3.74 | 6.41 | CV_C |
| AX.166519032 | 2C | 183.5 | 3.04 | 5.07 | STR |
| **AX.166521303** | **2B** | **63.1** | **3.29** | **9.41** | **CIR** |
| AX.89791395 | 4B | 130.4 | 3.17 | 2.71 | CIR |
| AX.166509340 | 4C | 241.7 | 3.00 | 2.37 | CIR |
| AX.89890707 | 5C | 212.2 | 3.11 | 1.86 | CIR |
| AX.166526395 | 7C | 228.6 | 3.81 | 3.95 | CIR |
| AX.89873861 | 1A | 139.7 | 3.73 | 5.27 | L/W |
| AX.123357666 | 3B | 303.6 | 3.35 | 3.42 | L/W |
| AX.89829301 | 4D | 182.3 | 3.03 | 2.06 | L/W |
| AX.166524494 | 5B | 103.6 | 3.25 | 3.83 | L/W |
| **AX.89791880** | **5D** | **123.7** | **3.52** | **1.46** | **L/W** |
| AX.89792701 | 5D | 182.0 | 3.61 | 10.64 | L/W |
| AX.166503621 | 2C | 266.2 | 3.89 | 2.19 | CV_D |
| AX.166504095 | 3A | 149.5 | 4.10 | 2.8 | CV_D |
| AX.166525754 | 6C | 74.7 | 3.25 | 3.58 | CV_D |
| **AX.166521303** | **2B** | **63.1** | **4.55** | **15.08** | **Max_A/Min_A** |
| **AX.89791880** | **5D** | **123.7** | **4.22** | **3.11** | **Max_A/Min_A** |
| AX.166506309 | 5D | 150.8 | 3.16 | 2.48 | Max_A/Min_A |
| AX.166523943 | 5D | 233.9 | 3.42 | 4.47 | Max_A/Min_A |
| **AX.166521303** | **2B** | **63.1** | **3.86** | **6.28** | **CV_A** |
| AX.89788547 | 5D | 40.9 | 3.61 | 5.8 | CV_A |

**Supplementary Figure 1.**
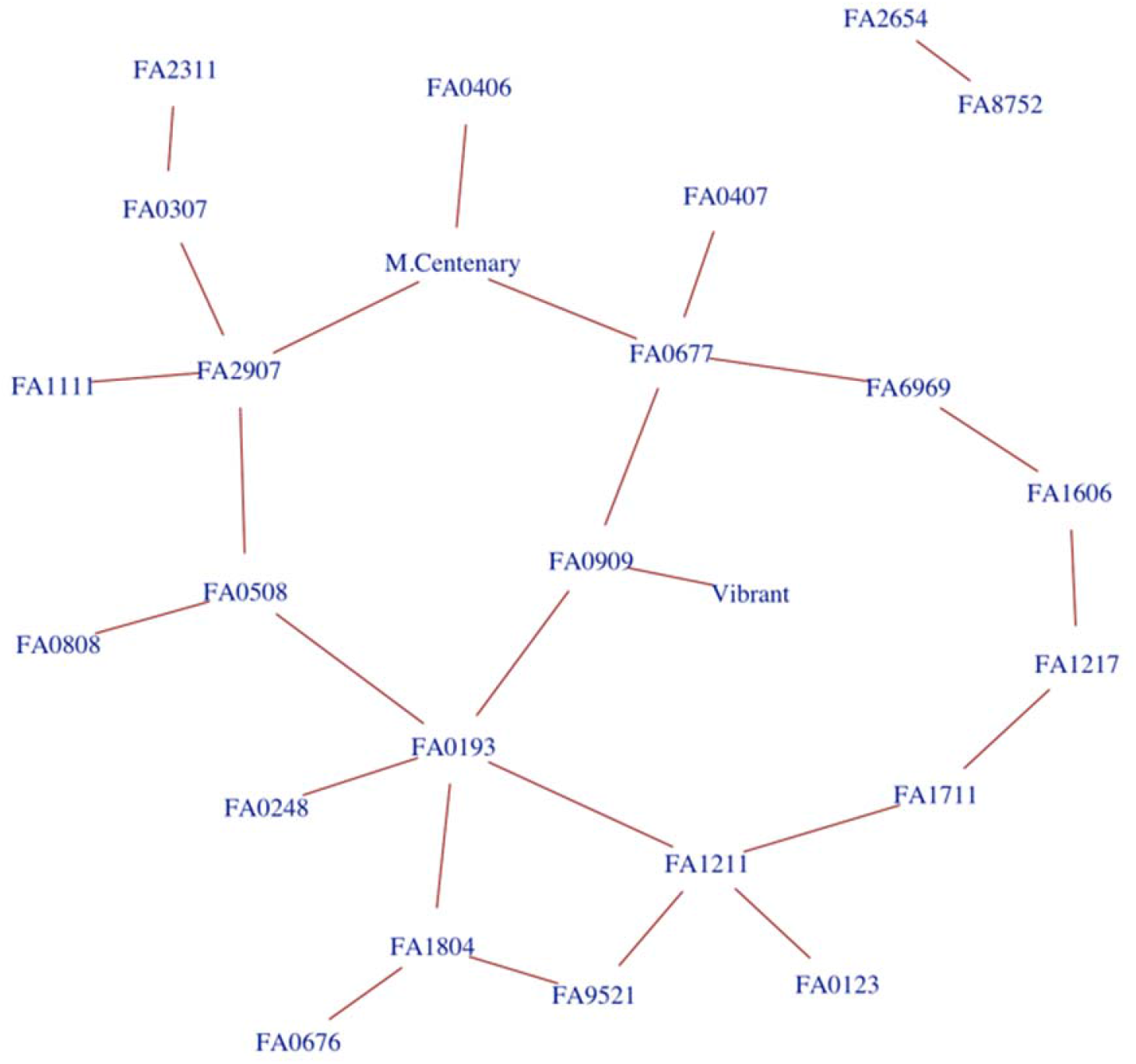
A network of crosses conducted to generate the multiparental mapping population used in this study. Cultivars are represented by circles, families are represented by lines.

**Supplementary Figure 2.**
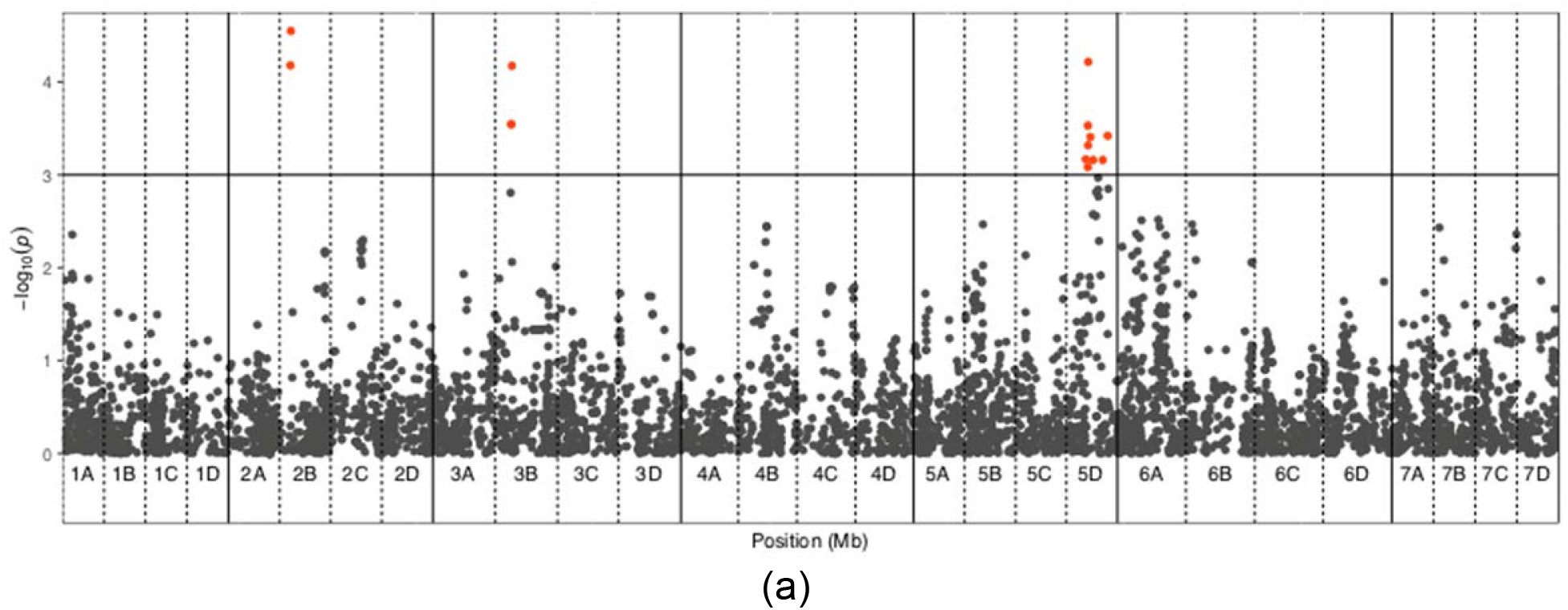
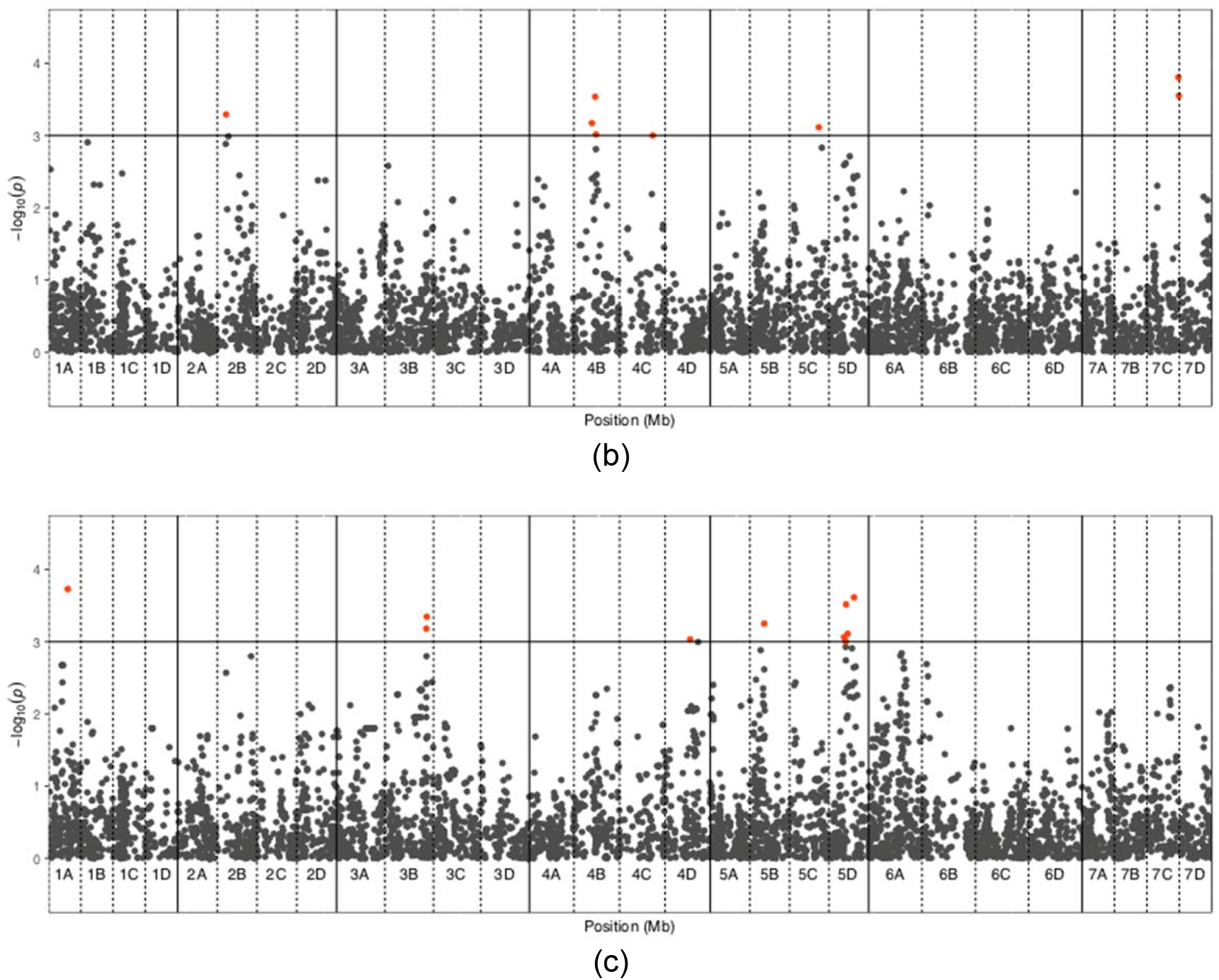
Manhattan plots of (a) Max_A/Min_A, (b) L/W and (c) CIR.

